# Leapfrog dynamics in phage-bacteria coevolution revealed by joint analysis of cross-infection phenotypes and whole genome sequencing

**DOI:** 10.1101/2020.10.31.337758

**Authors:** Animesh Gupta, Shengyun Peng, Chung Yin Leung, Joshua M. Borin, Sarah J. Medina, Joshua S. Weitz, Justin R. Meyer

**Affiliations:** Department of Physics, University of California San Diego, La Jolla, CA, USA; School of Biological Sciences, Georgia Institute of Technology, Atlanta, GA, USA; Division of Biological Science, University of California San Diego, La Jolla, CA, USA; School of Physics, Georgia Institute of Technology, Atlanta, GA, USA

## Abstract

Viruses and their hosts can undergo coevolutionary arms races where hosts evolve increased resistance and viruses evolve counter-resistance. Given these arms race dynamics (ARD), viruses and hosts are each predicted to evolve along a single trajectory as more recently evolved genotypes replace their predecessors. Here, by coupling phenotypic and genomic analyses of coevolving populations of bacteriophage λ and *Escherichia coli*, we find conflicting evidence for ARD. Virus-host infection phenotypes fit the ARD model, yet whole genome analyses did not. Rather than coevolution unfolding along a single trajectory, cryptic genetic variation emerges during initial virus-host coevolution. This variation is maintained across generations and eventually supplants dominant lineages. These observations constitute what we term ‘leapfrog’ coevolutionary dynamics, revealing weaknesses in the predictive power of standard coevolutionary models. The findings shed light on the mechanisms that structure coevolving ecological networks and reveal the limits of using phenotypic assays alone in characterizing coevolutionary dynamics.

## Main text

### Introduction

Bacteria and their viruses (phage) are the two most abundant and genetically diverse groups of organisms on Earth (Torsvik *et al*. 2002; Clokie *et al*. 2011; Thomas *et al*. 2011). Together, bacteria and phage are part of ecological communities characterized by complex networks of interactions whose structures have important implications that extend beyond the microbial world. For example, when viral lysis of bacteria redirects organic matter towards the microbial loop and away from higher tropic levels, potentially reducing productivity of macroscopic organisms (Fuhrman 1999; Wilhelm & Suttle 1999; Suttle 2007; Weitz & Wilhelm 2012; Brum *et al*. 2016). Changes in phage-bacteria infection networks that arise due to evolution of resistance (or counter-resistance) may have ripple effects throughout ecological communities and associated ecosystems.

Viral infection and lysis represents a strong selective pressure for the evolution of phage resistance (e.g., (Luria & Delbrück 1943) and the corresponding evolution of expanded host ranges amongst phage (Luria 1945). Evolutionary changes in resistance and infectivity lead to dynamic arms races where bacteria evolve phage resistance (Labrie *et al*. 2010), and phage evolve counter defenses (Hampton *et al*. 2020). This dynamic causes continual remodeling of the interaction network, which can impact ecosystem processes, the stability of ecological communities, and maintenance of microbial diversity (Bohannan & Lenski 2000; Buckling & Rainey 2002; Rodriguez-Valera *et al*. 2009; Stern & Sorek 2011). There is a growing interest in characterizing the dynamics of phage-bacteria coevolution and to understand the molecular and ecological mechanisms that shape their coevolving networks (Koskella & Brockhurst 2014).

A starting point to study phage-bacteria coevolution is to characterize how their interactions change over time (Weitz *et al*. 2005; Childs *et al*. 2012; Weinberger *et al*. 2012; Valverde *et al*. 2017). Models of coevolution tend to predict two types of dynamics (Agrawal & Lively 2002; Weitz *et al*. 2013). One such model is that of arms race dynamics (ARD) where bacteria evolve resistance to an increasing number of phage and phage counter by expanding their host range. For the bacteria, this leads to an escalation where increasingly resistant bacteria replace their less-resistant predecessors. The increase of resistance is expected to cause rapid bacterial genomic divergence and an imbalanced phylogenetic pattern with a single pronounced branch. Similarly, as the phage broadens its host range, the most recently evolved type is expected to supplant its predecessors, resulting in the formation of a similarly imbalanced phylogeny. A second model of coevolution is based on the evolution of specialized interactions, often described as *lock and key* interactions (Weitz *et al*. 2013). In this model, as coevolution progresses, bacteria gain resistance to contemporary phage, but lose resistance to phage encountered in the past. Likewise, as phage evolve counter-defenses, they lose the ability to infect other host genotypes. Under this model, host genotypes rise and fall according to how abundant their corresponding parasite genotypes are, while parasite genotypes track the abundance of their hosts creating a feedback loop and fluctuating selection dynamics (FSD) (Sasaki 2000; Gandon *et al*. 2008). FSD produces negative frequency-dependent selection that is expected to promote diversification and the formation of a balanced phylogeny with multiple branches. ARD and FSD represent distinct archetypes of possible coevolutionary dynamics and models have been proposed that span the space between the two end points (Leung & Weitz 2016).

One way to gain insight on whether phage and bacteria coevolve according to ARD or FSD is to quantify their interaction networks (phage-bacteria interaction networks; PBINs) and test for nonrandom nested and modular patterns (Flores *et al*. 2011). Nestedness measures the extent to which interaction patterns form strict hierarchical subsets, analogous to nesting Russian dolls (Almeida-Neto *et al*. 2008). This pattern can be produced by ARD since at each step the phage adds on to its existing host range, expanding its range of infectable hosts in a way that encapsulates its ancestors’ ranges; likewise bacteria evolve resistance to previously evolved phage encapsulating its ancestors’ range of phage to which it is resistant. Modularity arises in networks when groups of phage and bacteria tend to interact significantly more often within clusters than between clusters. This pattern is consistent with FSD where interactions are expected to be highly specialized. The majority of PBINs are significantly nested supporting the prominence of ARD; however some PBINs are modular (Flores *et al*. 2011), and while rare, specialized interactions have been documented to evolve during phage-bacteria coevolution (Gómez & Buckling 2011). Surprisingly, nested patterns at short spatial scales can give way to modular patterns at large spatial scales (Flores *et al*. 2013), and ARD has been shown to give way to FSD during advanced stages of coevolution (Hall *et al*. 2011; Flores *et al*. 2013). Together, this variation and scale-dependence provide a glimpse of the challenges in connecting coevolutionary process to phenotypic outcome.

While the phenotypic predictions for ARD and FSD are often tested, assessments of the phylogenomic predictions are not as common (ARD: imbalanced, FSD: balanced), and we are unaware of an example where PBINs have been coupled with phylogenomic analyses (note that the link between evolutionary interactions and phylogenomic structure is relatively well developed in studies of virus infection of human and animal hosts, sensu (Koelle *et al*. 2006; Volz *et al*. 2013)). Here, we utilized a model phage-bacteria coevolutionary system; bacteriophage λ and its host, *Escherichia coli*. When these species are cultured under certain laboratory conditions, they rapidly coevolve (Meyer *et al*. 2012; Meyer & Lenski 2020). *E. coli* is known to evolve resistance through mutations in the regulatory gene *malT* that suppress expression of the host receptor, the outer-membrane protein LamB. λ counters this by evolving mutations in the binding domain of its host recognition protein J that allows it to use a new receptor, OmpF. *E. coli* then evolves additional mutations in OmpF or in an inner-membrane protein complex, ManXYZ, that transports λ DNA into the cytoplasm (Erni *et al*. 1987; Esquinas-Rychen & Erni 2001). While much is already known about the molecular details of their coevolution, little is known on the joint changes in interaction networks and genetic relatedness.

For this study, we revived cryopreserved samples that were isolated from a previously reported coevolution experiment (Meyer *et al*. 2012). We focused our analyses on a single replicate; the first experimental community in which λ evolved to use OmpF. We isolated a total of 50 bacteria and 44 phage spread across multiple time points. Next, we constructed a PBIN of all combinations of pairwise phage and bacteria interactions and used multiple analyses to characterize their coevolution based on phenotypes. All three phenotype-based analyses suggested that viruses and microbes engage in ARD. In parallel, we sequenced the full genomes of each isolate and reconstructed the isolates’ phylogenetic relationships. The genome sequences revealed a phylogenetic pattern that was inconsistent with the ARD model. Our study demonstrates that phenotypic analyses are not sufficient to test hypothesis on coevolutionary dynamics and reveals a new type of coevolutionary dynamic we refer to as Leapfrog Dynamics (LFD).

## Materials and methods

### Details on the initial coevolution experiment previously published

Meyer *et al*. (Meyer *et al*. 2012) performed the original coevolution experiment with *Escherichia coli* B strain REL606 and a lytic bacteriophage λ strain, cI26. This λ strain was chosen because it cannot initiate lysogeny, a life cycle phase where λ confers immunity to additional λ infections. By choosing a lytic strain, we forced the bacteria to evolve genetic resistance. *E. coli* and λ were cocultured in a carbon-limited minimal glucose media at 37 °C for 37 days (Meyer *et al*. 2012). At the end of each day, 1% of the community was transferred to new flasks with fresh media, and, weekly, 2 ml of the community was preserved by adding ~15% of glycerol and freezing the mixture at −80 °C.

### Isolation of host and phage clones

We randomly isolated ten host and eleven phage individuals from different timepoints from the cryopreserved samples. In total, 50 strains of *E*. coli and 44 strains of λ were isolated from days 8, 15, 22, 28 and 37 of the experiment (no phage were detected on day 37). Bacteria were isolated by streaking onto Luria-Bertani (LB) agar plates (Sambrook & Russell 2001) and randomly picking 10 colonies. These colonies were re-streaked three times to remove phage particles and grown overnight in liquid LB to create stocks. Phage were isolated by plating an appropriate dilution of the population onto overlay plates (Adams 1959) with the sensitive ancestral bacteria, REL606, and randomly picking 11 plaques. These plaques were grown overnight with REL606 in LBM9 medium and stocks were created using chloroform isolation technique (Meyer *et al*. 2012). All phage and bacteria stocks were stored at −80 °C with the addition of 15% *v/v* glycerol.

### Pairwise infection assays and efficiency of plating (EOP)

We performed quantitative, pairwise infection assays for all combinations of host strains and phage strains that were isolated. Specifically, seven serial 1/10^th^ dilutions were made of each phage isolate. 2 μl of each dilution plus undiluted phage stock was spotted on top of different host strain lawns including ancestor REL606. Thus, a total of 8*44 spots of phage were plated on 51 different types of bacterial lawns, leading to a total of 17,952 pairwise infections. Pairwise infection experiments measure how well each phage isolate infects every host isolate. We quantified phage infectivity by calculating efficiency of plating (EOP), defined as the ratio of density of phage isolate calculated on a coevolved isolate to the density of phage calculated on the REL606 ancestor.

### Analysis of PBIN nestedness and modularity

The *BiMat* software was used to assess the nestedness and modularity of the PBIN (Flores *et al*. 2016). The raw EOP value matrix was binarized into 0 for EOP = 0 and 1 for EOP > 0, and then *BiMat* was run with default settings. Here we report the statistics for a conservative version of the analysis where the rows and columns that contained all zeros were removed from the matrix to reduce any bias these entries cause in establishing significant nested patterns.

### Resistance and infectivity calculations

For a total number of *n* host samples and *m* phage samples, we denote the EOP value for the *i*th host sample against *j*th phage sample as *e_ij_* where *i* ∈ [1, *n*] and *j* ∈ [1, *m*]; *n* = 50 and *m* = 44. We denote the five checkpoint days of day 8, 15, 22, 28 and 37 for host by *k*, where *k* = 1,2,3,4,5, and the four checkpoint days of day 8, 15, 22 and 28 for phage by *l* where *l* = 1,2,3,4. Host resistance for a host sample *i* is calculated as

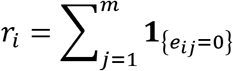

where **1**_*X*_ is the indicator function and *r_i_* measures the number of phage strains that the host is resistant to. The host range of a phage sample *j* is calculated as

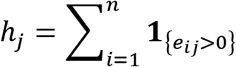

which measures the number of host strains that the phage can successfully infect. The resistance percentage of host for each day is calculated as

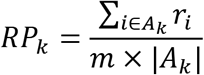

where *A_k_* denotes the range of the host sample that belongs to the *k*th sample checkpoint and |*A_k_*| denotes the cardinality of the set *A_k_*, i.e., the number of host samples at the *k*th checkpoint. Likewise, the host range percentage of phage for each day is calculated as

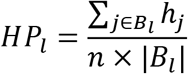

where *B_l_* denotes the range of the phage sample that belongs to the *l*th checkpoint and |*B_l_*| denotes the cardinality of the set *B_l_*, i.e., the number of phage samples at the *l*th checkpoint.

### Genomic DNA preparation for sequencing

λ genome extraction for whole genome sequencing were previously reported in (Meyer *et al*. 2016). To summarize, λ particles were concentrated using PEG precipitation, the phage were treated with DNase I to remove free-floating DNA not protected by phage capsids, the DNase is denatured with heat, which also releases capsid-enclosed phage DNA. The DNA was extracted using Invitrogen’s PureLink kit. *E. coli* genomic DNA was extracted and purified from a 1 ml sample of culture by using PureLink kit. Genomic DNA was further processed by fragmenting the DNA and attaching adapters and barcodes using a method outlined in (Baym *et al*. 2015). Sequencing was done at UC San Diego IGM Genomics using paired-end Illumina HiSeq 4000 platform.

### Construction of mutation profile tables

After collecting the raw sequencing reads, we removed the adapters using cutadapt (Martin 2011) and performed quality control (QC) for each isolated strain using FastQC (Andrews 2010). The QC filtered sequencing reads were then analyzed using the *breseq* (v0.32.1) (Deatherage & Barrick 2014). We ran *breseq* in the consensus mode with default parameters except for the consensus-frequency-cutoff, which was set to 0.5.

### Phylogenomics

Due to the prevalence of large insertions and deletions in the host genomes, conventional nucleotide substitution models were not suitable for estimating the host phylogenetic tree. However, such models were suitable for estimating the maximum-likelihood phylogenetic tree for phage genomes. As a result, two different approaches were taken to reconstruct the evolutionary trajectories of the host and virus.

To construct the phage phylogeny, multiple sequence alignments were performed for all recovered genomes and the ancestral genome using *mafft* (v7.305b) (Katoh *et al*. 2002) with default settings except that ‘retree’ was set to 2 and ‘maxiterate’ was set to 1,000. A maximum likelihood tree was constructed using *raxml-ng* (Stamatakis 2014). Finally, the *TreeTime* (Sagulenko *et al*. 2018) program was used to generate the phylogenetic tree.

To reconstruct the host’s phylogeny, we constructed a Hamming distance matrix to calculate genetic distances between different host isolates. Neighbor-joining (NJ) trees were then built based on the hamming distance matrix using *T-REX* (Makarenkov 2001). Finally, the *TreeTime* program was used to build the host phylogenetic tree.

### Whole genome whole population sequencing

We sequenced the full population of phage and bacteria from Day 8 of the experiment to 3,726-fold and 142-fold coverage, respectively. Whole population sequencing uncovered alleles in the bacterial and phage populations that existed at lower frequencies than we could detect by isolating individuals. To do this, λ and *E. coli* populations were revived by growing 120 μl of frozen stock of the whole community in the laboratory conditions from the original experiment (Meyer *et al*. 2012). Phage and bacteria were then separated, and their genomic DNA was extracted in the same manner as described before for clonal stocks. Genomic libraries were prepared using NexteraXT kit at UC San Diego IGM Genomics. IGM also sequenced the samples using 75 base single reads on the Illumina HiSeq 4000 platform. *breseq* v0.32.1 was used to analyze whole population sequencing data of Day 8. We ran *breseq* in polymorphism mode with default settings to construct the mutation profile tables.

## Results

### Phage-bacteria infection network

The pairwise interaction study revealed multiple λ genotypes with phenotypically distinct host ranges, and *E. coli* genotypes that vary in resistance (Fig. 1a). In line with the ARD model, we found that the interactions were highly nested (Fig. 1b) and had a low level of modularity (Fig. S1). Also, in line with ARD, *E. coli* evolved increasing resistance (Fig. 1c, 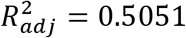, *F*_1,48_ = 51.01, *P* = 4.453e-09 for linear model: response ~ time), and λ gained increasing host range and infectivity (Fig. 1d, 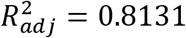, *F*_1,42_ = 188.1, *P* = 2.2e-16) through time. Note, all mean EOP values were zero for day 8 phage because all isolated hosts were resistant to all day 8 phage (Fig. 1a and 2a).

**Fig. 1.**
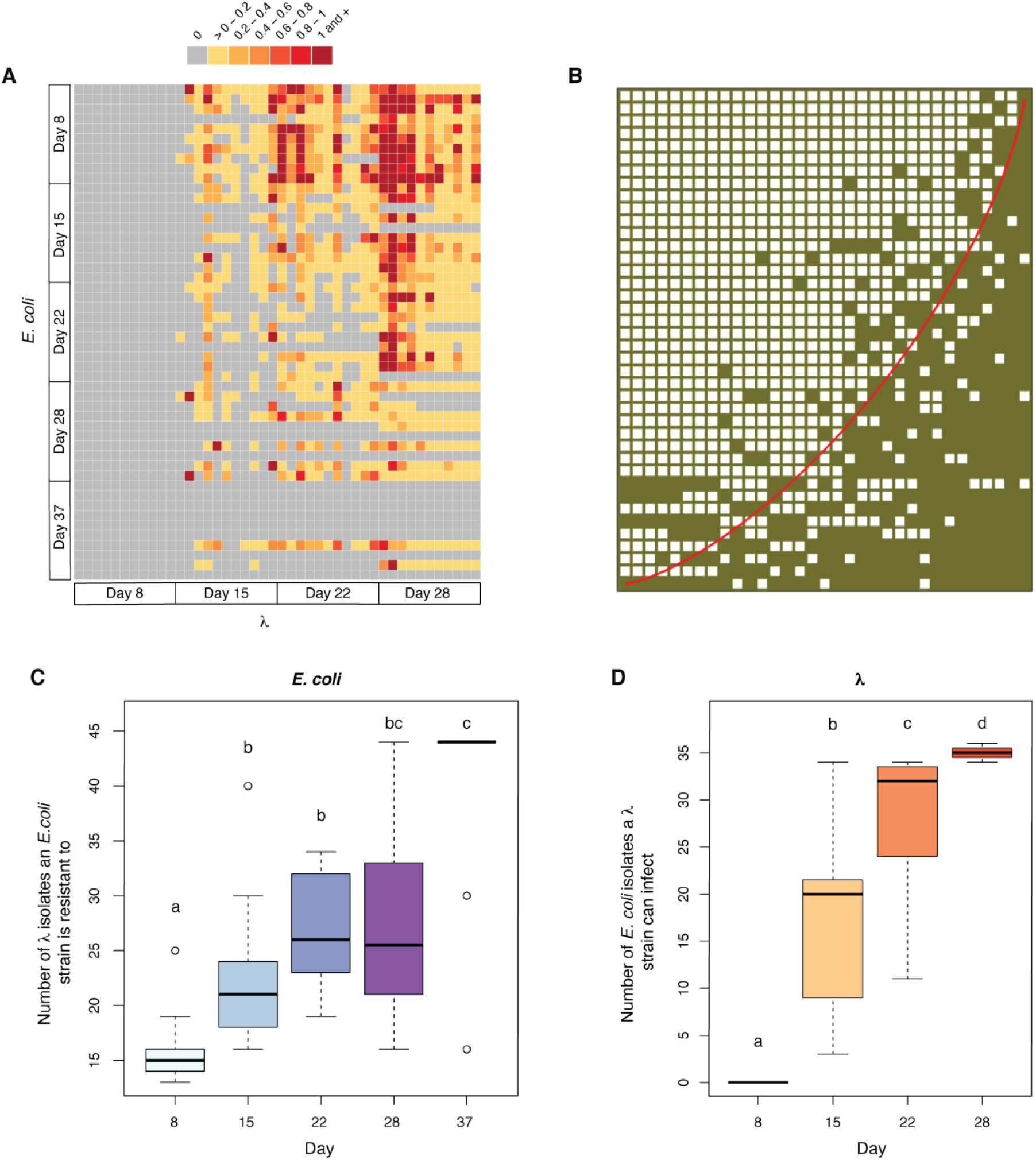
Host resistance and phage infectivity measured by pairwise plaque assays. (**A)** Phage-bacteria infection network where the color of each cell is determined by the EOP values obtained for that host-phage interaction pair; grey cells represent no infection by λ on the given *E. coli* strain, yellow represents low infectivity and red represents high infectivity. (**B)** The original network in a) reassembled using the software *BiMat* to visualize maximal nestedness (Flores et al., 2016). Filled squares indicate a combination of host and phage that result in successful interactions (EOP > 0), and the red line highlights the isocline using the nestedness temperature calculator (NTC) algorithm. The nestedness value of the network utilizes the nestedness based on overlapping and decreasing fill (NODF) metric, which was significantly greater than the null expectation when constraining the fill of the bipartite network (measured value of nestedness 0.839 vs. null value of nestedness 0.638 ± 0.011 based on 200 trials). **(C)** Boxplots showing the total number of λ isolates from all days that *E. coli* genotypes are resistant to across different sampling days. **(D)** Boxplots showing the total number of *E. coli* isolates from all days that λ genotypes can infect across different sampling days. Lowercase letters in c) and d) denote significant difference between different days via Tukey’s honest significance test: c) ANOVA: *F*_4,45_ = 13.3, *P* = 3.11e-07), d) ANOVA: *F*_3,40_ = 67.05, *P* = 1.17e-15. A simple linear regression model with time as the predictor variable was also used to test if *E. coli* evolved increasing resistance in c) and λ evolved increasing host range in d) (statistics in the main text).

**Fig. 2.**
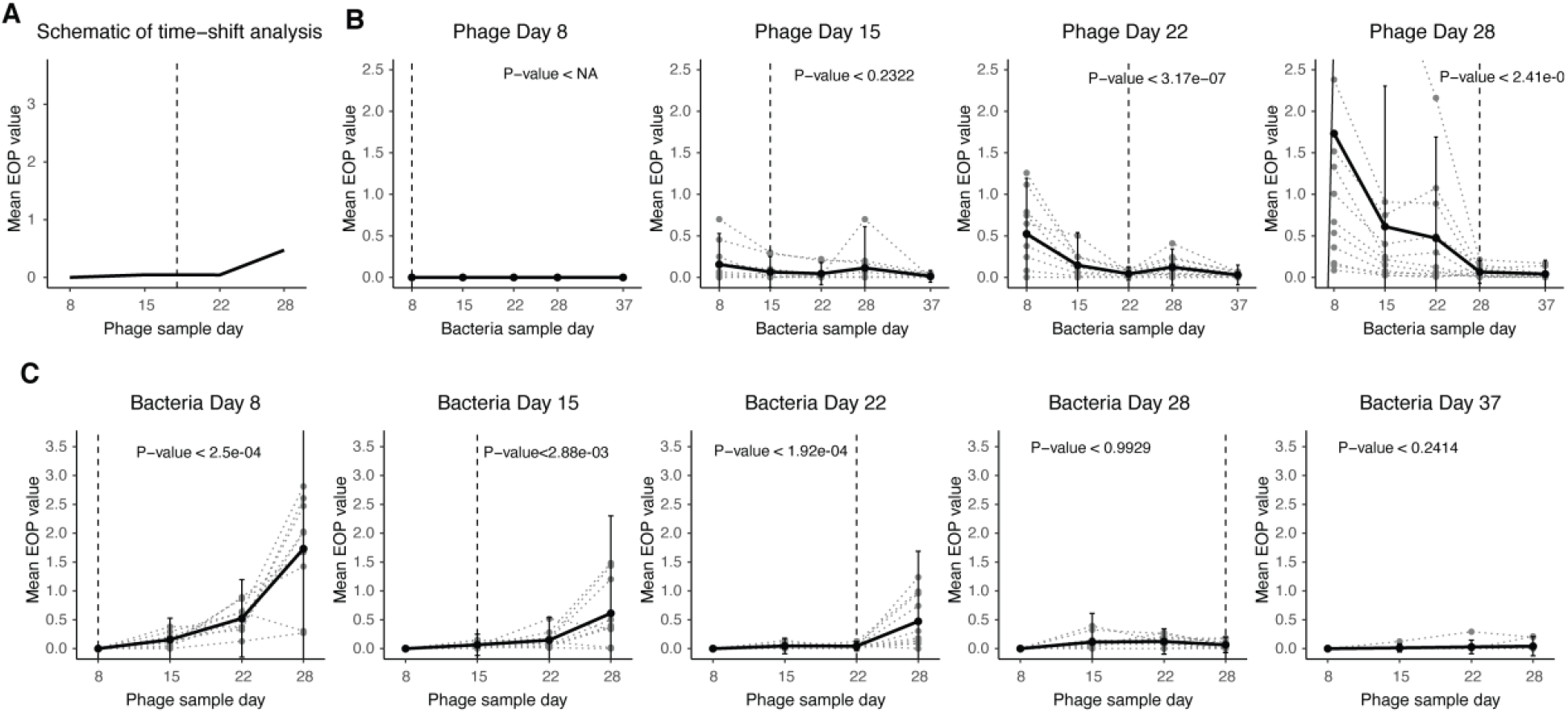
Time-shift analysis results from different checkpoints. (**A)** Schematic for the time-shift analysis that compares the mean EOP from hosts or phage interacting with their counterparts from the past, contemporary and the future. **(B)** Time-shift results from phage checkpoints day 8, 15, 22 and 28 respectively. The gray dotted line shows the time-shift curve for each individual phage and the black line shows the average. The vertical dashed line represents the phage sample day. The *P*-values shown here are the maximum *P*-value from one-sided paired *t*-tests comparing the initial checkpoints with each of the later checkpoints. **(C)** Time-shift results from host checkpoints day 8, 15, 22, 28 and 37, respectively. The gray dotted line shows the time-shift curve for each individual host and the black line shows the average. The vertical dashed line represents the host sample day. The *P*-values shown here are the maximum *P*-value from one-sided paired *t*-tests comparing the final checkpoints with each of the previous checkpoints.

To further test between indicators of ARD and FSD, we performed a time-shift analysis using efficiency of plating (EOP) values to determine how phage infectivity varies when presented with past, contemporary, or future bacteria (Fig. 2a) (Gaba & Ebert 2009). ARD predicts that phage will be able to infect past and contemporary, but not future hosts, while FSD predicts that phage will be best at infecting contemporary hosts. The time-shift analysis was conducted for each λ isolate by calculating its mean EOP value for all 10 bacterial isolates on each day (Fig. 2b). This analysis was repeated for the bacteria using the same EOP data but by calculating levels of resistance to λ isolates from different time points (Fig. 2c). In line with ARD, λ isolates from days 22 and 28 had higher infectivity on past hosts than contemporary or future hosts (Fig. 2b). The analysis for day 15 phage was inconclusive because the EOP values across time were not statistically significant. The pattern for bacteria was also in line with ARD: isolates from days 8, 15 and 22, had lower resistance (higher EOP) for phage samples from the future versus the phage isolated from the same time or in the past (Fig. 2c). A full time-shift analysis could not be conducted for isolates from days 28 and 37 since the phage went extinct between days 28 and 37, however isolates from these time points were the most resistant.

At all timepoints, phage were isolated using the ancestral host (REL606) in an effort to minimize sampling bias. Notably, in previous studies we found that all evolved phages were able to infect the ancestor (Flores *et al*. 2011). However, if phages evolve to specialize on coevolved bacterial genotypes in line with FSD, they may have lost the ability to infect the ancestor and not be sampled, thereby artificially reducing support for FSD and strengthen the signal of ARD. To test whether phages evolved that lost the ability to infect REL606, we isolated phages using lawns of all unique *E. coli* genotypes (16 unique genotypes, however no plaques formed on 3). Eight phages were sampled from each host yielding a sample size of 104 phage isolates. Each phage isolate was able to form plaques on REL606 (Table S3, Supplementary Information). An additional efficiency of plaquing analyses was performed for a subset of the phages; one phage isolate from each host. This analysis revealed that most phages were more likely to produce plaques on REL606 than on the coevolved hosts from which they were isolated (Table S3, Supplementary Information). Together, these results suggest that REL606 was suitable for sampling the coevolved phage diversity in this system.

### Genome sequencing

The genome sequencing revealed 22 unique *E. coli* genomes and 34 unique phage genomes. Among the *E. coli* strains, we found a total of 18 unique mutations: 6 missense mutations, 1 nonsense mutation, 1 intergenic point mutation, 7 deletions and 3 duplications (Fig. 3a). The most abundant mutation that occurred in 38 out of 50 host genomes was a frameshift mutation caused by a 25-base duplication in the *malT* gene, in line with the original study (Meyer *et al*. 2012). Disruptions in *malT* interferes with the expression of LamB protein which ancestral λ needs to bind to *E. coli* cells. We also observed one isolate with a *lamB* mutation (1-base deletion) *in lieu* of the typical *malT* mutation. The most resistant *E. coli* strains on day 37 have multiple mutations that are expected to confer resistance; a *malT* deletion, a nonsynonymous change in *ompF*, and a deletion in *manZ* (Meyer *et al*. 2012; Burmeister *et al*. 2021).

**Fig. 3.**
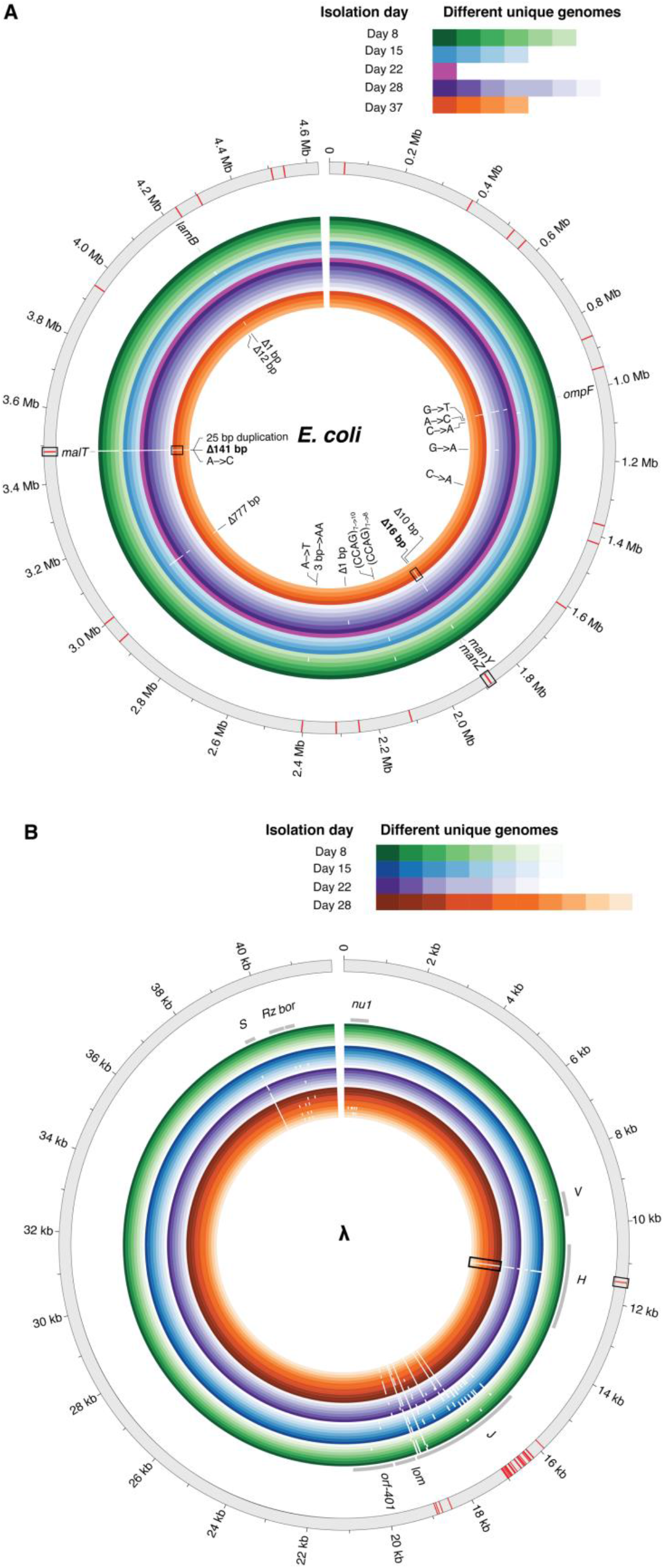
Genomic diversity in clones isolated from different days and full population sequencing for (A) *E. coli* and (B) λ. The outermost gray ring represents the reference genome with red bars indicating the placement of mutations uncovered by the whole-population sequencing at day 8. The inner colored rings represent the isolates sequenced from different time points (outer rings are genomes isolated from earlier time points). Shades within each color depict unique genomes sequenced from each time point. White gaps in the genomic rings indicate the location of mutations. All mutations found in clonal isolates have been labeled for *E. coli* in A); however, due to the large number of mutations in λ, only the gene names that harbor mutations have been identified (gray bars). The mutations that become dominant at later stages of coevolution and were also found in day 8 population sequencing have been highlighted with rectangular boxes.

One of the most common bacterial mutations observed in our dataset was a 777 bp deletion caused by the excision of an IS element. Despite occurring in 25 genomes, we had not observed the deletion in any previous studies and the deletion did not affect any genes known to protect against λ infection; *insB-22* which encodes for IS1 protein InsB, *insA-22* which encodes for IS1 protein InsA, and *ECB_02825* which encodes for a pyrophosphorylase (Maynard *et al*. 2010). Similar IS mutations are known to occur at high rates (Cooper *et al*. 2001) and the deletion was only observed in genomes with a *malT* mutation, suggesting it might be a neutral genomic hitchhiker. We tested this hypothesis in a follow-up experiment which revealed that the deletion enhances resistance when it cooccurs with the *malT* mutation, suggesting the deletion was adaptive and epistatic with *malT* mutations (Fig. S2, Supplementary Information). We are unsure of how the deletion confers resistance; however, our results suggest that experimental evolution is likely to reveal novel sites in the λ-*E. coli* interactome than previously known.

In the λ genomes, we found a total of 176 unique mutations: 53 nonsynonymous point mutations, 87 synonymous point mutations, 2 insertions, 3 deletions and 31 intergenic mutations (Fig. 3b). While this level of molecular evolution may seem surprising for such a short-term experiment, similar levels have been observed for other phage evolving in the laboratory (Wichman *et al*. 2005). 116 of these mutations were in the host-recognition gene *J*. The J protein is positioned at the end of the phage’s tail, and initiates infection by binding to *E. coli*’s LamB protein. Some of these J mutations have been shown to increase adsorption rates to LamB and allow λ to exploit a novel receptor, OmpF (Burmeister *et al*. 2016; Maddamsetti *et al*. 2018; Petrie *et al*. 2018). Interestingly, the extensive sequencing effort performed here revealed a mutation in another tail fiber protein called H (C→T substitution at nucleotide position 11,451). This mutation rises late in the experiment and likely plays a role in expanding λ’s host range. The tape-measure gene *H* helps determine the length of λ’s tail, and mutations in this gene have been shown to increase λ’s host range (Scandella & Arber 1976).

### Phylogenomic reconstruction of coevolution

Even though multiple analysis of the phenotypic data supported the ARD model for coevolution, the pattern produced by the phylogenies are in line with predictions of FSD (Fig. 4). The phylogenies of both *E. coli* and λ show that multiple lineages coexist for weeks, rather than a single dominant branch. A second unexpected observation was that the bacteria that had acquired the highest level of resistance at the end of the experiment on day 37 was not most closely related to isolates at previous time points (e.g., days 28, 22, or 15), instead it was most closely related to a common ancestor of isolates identified at the early stages of the experiment (on day 8). This finding suggests that a rare lineage leapt ahead of the dominant lineage, a process we term leapfrog dynamics (LFD). Similarly, for λ, we found that that the clade dominant at the final timepoint with the broadest host range was more closely related to wildtype λ than the clade dominant at preceding timepoints. For both species, the clades that win out later in the arms race appeared to persist as cryptic subpopulations early in the coevolutionary experiment.

**Fig. 4.**
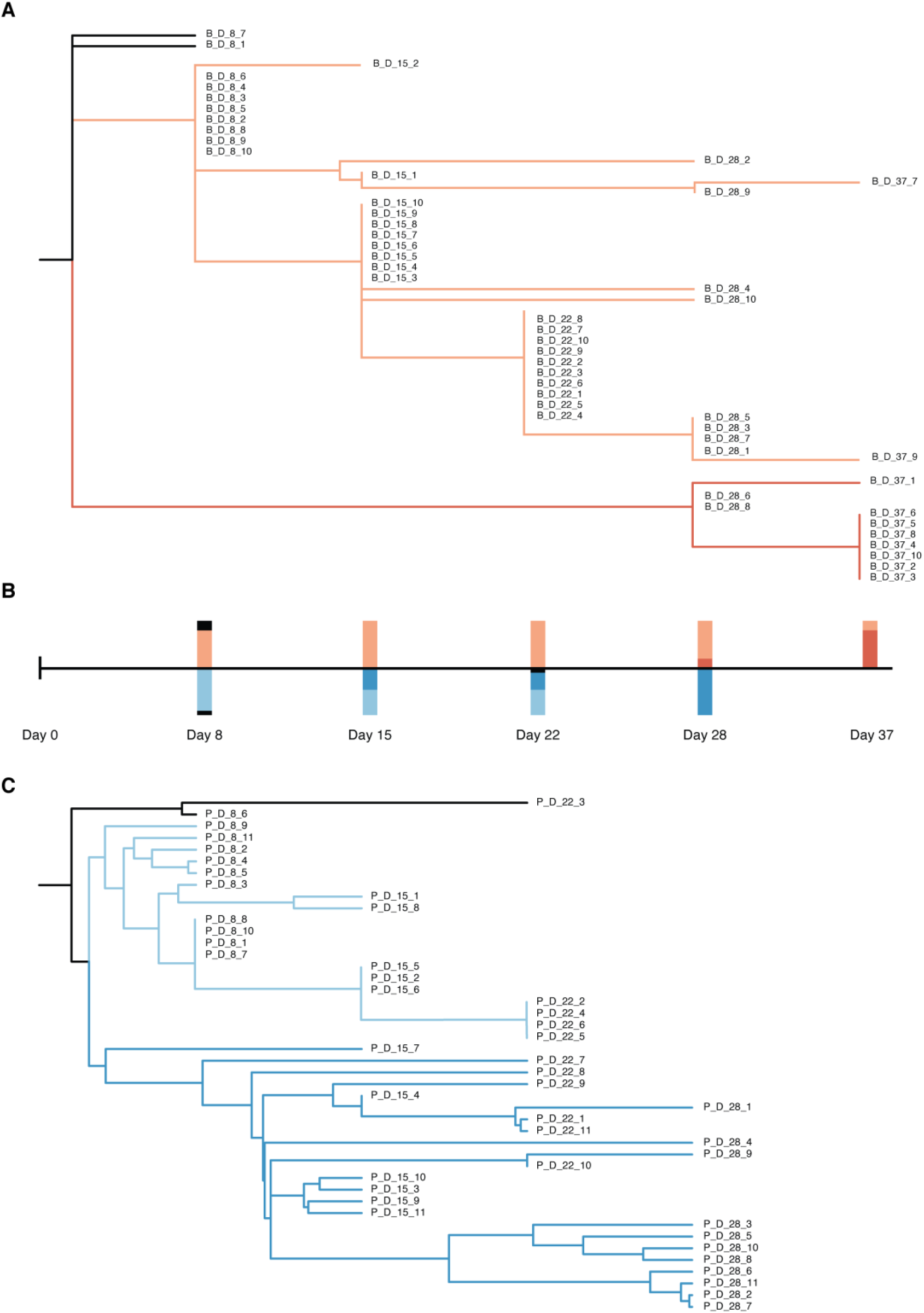
Reconstructed phylogenetic trees of the host and phage. **(A)** The host phylogenetic tree based on host mutation profiles. All completely-resistant host strains are located on the red branch. Bars above the time scale in (**B**) represents the proportion of host strains from each colored branch across different checkpoints. **(C)** The phage phylogenetic tree based on the phage mutation profiles. All day 28 phage strains are located on the dark blue branch. Bars below the time scale in (**B**) represents the proportion of phage strains from each colored branch across different checkpoints.

### Whole population sequencing at an early stage of coevolution

To test the key prediction of LFD that cryptic lineages coexist with dominant lineages and can supply the genetic reservoir used for later stages of coevolution, we sequenced full populations of *E. coli* and λ from day 8 and searched for mutations that rose to prominence at the end of the study. For *E. coli*, we were specifically searching for two mutations: a Δ16 bp deletion at position 1,882,915 in *manZ* and a non-synonymous mutation at 1,003,271 in *ompF*. These mutations are present in most of the day 37 isolates and are thought to confer resistance. The Δ16 bp deletion in *manZ* was detected, but not the *ompF* mutation (Table S1). We also found a 141 bp deletion in *malT* that cooccurs in the day 37 genomes with the *manZ* mutation. The *malT* deletion was at the same frequency, suggesting that these mutations were indeed linked and that they evolved sometime before day 8. For λ, we focused on the mutation in the tail fiber gene *H* that rises to dominance between days 22 and 28. Indeed, this specific H mutation was present at day 8 (Table S2). Unlike *E. coli*, we did not find any other mutations present in the day 28 isolates, suggesting that the *H* mutation was the first adaptation to occur in this lineage.

Besides revealing the eventual mutations associated with the winning lineages of the arms race, we also discovered much more genetic diversity via whole population sequencing than through isolate sequencing (as anticipated). Whole genome sequencing revealed 52 unique mutations in *E. coli* and 38 mutations in λ from full population sequencing compared to 7 and 30 through isolate sampling, respectively. The combined sequencing strategy suggests that there is significant genetic diversity generated at the earliest phases of the arms race that can become the grist for subsequent adaptation as the host (for phage) or phage (for hosts) change as a result of coevolution.

## Discussion

Through large scale phenotypic assays and whole genome sequencing, we were able to test existing paradigmatic models of coevolution and learn that each were inadequate to explain λ and *E. coli*’s coevolutionary dynamics. Three complimentary phenotypic analyses in Fig. 1 and Fig. 2 suggested that coevolution between λ and *E. coli* followed arms race dynamics (ARD). However, the phylogenetic pattern revealed by whole genome sequencing was consistent with fluctuating selection dynamics (FSD) (Fig. 3). These observations lead us to develop a new conceptual model to characterize λ and *E. coli*’s coevolution: which we term leapfrog dynamics (LFD). In this model, selection operates similarly to the ARD model, where parasite genotypes with ever-expanding host ranges are selected and hosts with ever-increasing resistance are favored. However, the difference is that in the LFD model there is a genetically diverse pool of hosts and parasites that evolve early, and on occasion, rare individuals are drawn from this pool with advantageous phenotypes and replace the dominant strains.

ARD models fall short in making accurate predictions for the phylogenies likely because of simplifying assumptions about the genetics of host range expansion and resistance. Evolutionary models tend to assume that mutations have additive effects on phenotypes (as highlighted in (Weitz *et al*. 2013)). Applied to ARD, this would mean that the phage with the broadest host range is likely to expand its host range faster than lagging genotypes, and for bacteria, the strain with the greatest resistance is most likely to acquire the next level of resistance and outcompete other strains. This genetic architecture favors the evolution of directed phylogenies with one dominant branch. Instead, λ J mutations are known to possess high-order epistasis and are nonadditive (Meyer *et al*. 2016; Maddamsetti *et al*. 2018). This may allow rare lineages with certain combinations of mutations to suddenly leap ahead if new modifications have synergistic interactions with preexisting mutations. Non-additivity was also discovered in the bacteria in this study with respect to the interactions between *malT^−^* and the Δ777 mutations (Fig. S2). A second problem with evolutionary models is the assumption of small effect-size mutations since mutations like the *malT* mutation can cause nearly complete resistance to some λ phage (Fig. 1a). Acquiring such large effect mutations could also help lineages leap ahead. Lastly, these models typically do not incorporate recombination, which was observed within this phage population to cause sudden increases in fitness (Borin *et al*. 2020) and could contribute to the phage’s ability to leap ahead. In contrast, evidence of recombination was not observed in the bacterial genome sequences, consistent with earlier findings in which recombination is not known to occur in this strain of *E. coli* (Souza *et al*. 1997).

One question left unanswered by this study is how multiple lineages persisted in this population for long durations. We hypothesized that trade-offs between host range and other viral traits for phage, and resistance and competitive fitness for bacteria, could explain the evolution of genomic diversity. Trade-offs between host range and λ stability were previously observed (Petrie *et al*. 2018); however, we were unable to detect trade-offs in *E. coli* for phage resistance. Our study also comes with limitations. We sampled clones at weekly resolution rather than sampling the full community at higher resolution (e.g., daily), which limits opportunities to quantitatively assess the emergence and changes in the frequency and linkage of mutations (e.g., (Lang *et al*. 2013)) as well as the extent of potential trade-offs given the changing ecological context.

Our findings have important implications for understanding the dynamic mechanisms that structure nested PBINs. The nested pattern is ubiquitous in PBINs (Flores *et al*. 2011), as well as many other ecological networks (Bascompte *et al*. 2003; Guimarães *et al*. 2007; James *et al*. 2012), so it is important to understand the processes that produce this ordering. One hypothesis is that the structure is determined by the genetics of the interactions (as in the gene-for-gene mechanism of coevolution, see (Weitz *et al*. 2013)). An alternative hypothesis for the emergence of nestedness is ecological: nestedness emerges because of how the selection steadily shifts during an arms race to promote incremental increases in host range and resistance. The conventional wisdom for which process controls the arms races is the underlying genetics. This stems from a pervasive idea that genetic mutation and evolution happen more slowly than changes in ecology, so coevolutionary systems must be constrained by their access to genetic variation. The coevolution experiment studied here was initiated with small population size of isogenic stocks of λ and *E. coli*. This should have favored genetic control because all of the variation had to evolve *de novo*. However, significant and evolutionarily relevant genetic variation was generated at the earliest phase of the arms race that appeared to remain cryptic in an ecological sense for many generations, suggesting that the ecology controlled the dynamics, not the availability of genetic variation.

Lastly, our results provide a cautionary tale for over-interpreting phenotypic data based on phage-bacteria infection networks and/or phenotype-based time shift experiments alone. Our initial prediction before performing genomic analysis was that λ-*E. coli* coevolution under these laboratory conditions would fit the ARD model. However, this proved to be erroneous due to the large amount of cryptic genetic variation and its role in driving late stages of coevolution. Interestingly, this study also reveals a potential limitation of using PBIN data for making dynamical predictions. Based on the matrix in Fig. 1a one would predict that the phage should go extinct on day 8. Still, λ did not go extinct in the coevolution experiment, consistent with a phenomenon known as ‘leaky-resistance’ (Chaudhry *et al*. 2018). Leaky resistance denotes a phenomenon by which a small number of resistant hosts revert to sensitive, thereby sustaining phage in the population until phage evolve to gain access to a new receptor, e.g., OmpF in the case of phage λ (Meyer *et al*. 2012). Together, the mechanisms of LFD and leaky-resistance show that host-phage dynamics revealed from PBINs alone miss the rich dynamics occurring at lower frequencies in the population.

In summary, by studying coevolving phage and bacterial populations with both phenotypic and genomic approaches we revealed emergent coevolutionary patterns that are not wholly explained by archetypical models of coevolution. Moreover, we found that phenotypic assays alone fall short in characterizing the underlying nature of coevolution – given the potential for cryptic genetic variation to fundamentally alter coevolutionary trajectories. We also showed that highly organized ecological pattern like nestedness can emerge despite the apparent absence of fundamental genetic constraints, demonstrating the power of selection in driving emergent ecological patterns. Moving forward, these results suggest the critical need to incorporate both phenotypic and phylogenomic approaches in evaluating phage-bacteria coevolution.

## Acknowledgements

We would like to thank Morgan Mouchka for her help with sequencing genomes of the whole community.

## Author Contributions

JRM isolated the viruses, isolated the bacteria, and measured the phage-bacteria infection matrix. AG and CYL sequenced whole genomes. AG and SP performed analyses, AG, CYL, JMB and SJM performed experiments, AG, SP, JSW and JRM wrote the manuscript, and JRM and JSW developed the project and oversaw the research. All coauthors helped edit the manuscript.

## Data accessibility statement

All sequencing data will be deposited in the NCBI Sequence Read Archive. Code will be made available at GitHub and data that support the findings of this study will be deposited at Dryad. Biological material is available from J.R.M. under a material transfer agreement with UC San Diego (http://blink.ucsd.edu/research/conducting-research/mta/index.html).

## References

1. Adams, M.H. (1959). Bacteriophages. Interscience Publishers, Inc., New York.

2. Agrawal, A. & Lively, C.M. (2002). Infection genetics: gene-for-gene versus matching-alleles models and all points in between. Evolutionary Ecology Research, 4, 79–90.

3. Almeida-Neto, M., Guimarães, P., Guimarães Jr, P.R., Loyola, R.D. & Ulrich, W. (2008). A consistent metric for nestedness analysis in ecological systems: reconciling concept and measurement. Oikos, 117, 1227–1239.

4. Andrews, S. (2010). FastQC: a quality control tool for high throughput sequence data.

5. Bascompte, J., Jordano, P., Melián, C.J. & Olesen, J.M. (2003). The nested assembly of plant-animal mutualistic networks. Proc Natl Acad Sci U S A, 100, 9383–9387.

6. Baym, M., Kryazhimskiy, S., Lieberman, T.D., Chung, H., Desai, M.M. & Kishony, R. (2015). Inexpensive Multiplexed Library Preparation for Megabase-Sized Genomes (vol 10, e0128036, 2015). Plos One, 10, 1.

7. Bohannan, B.J.M. & Lenski, R.E. (2000). Linking genetic change to community evolution: insights from studies of bacteria and bacteriophage. Ecology Letters, 3, 362–377.

8. Borin, J.M., Avrani, S., Barrick, J.E., Petrie, K.L. & Meyer, J.R. (2020). Coevolutionary phage training leads to greater bacterial suppression and delays the evolution of phage resistance. bioRxiv, 2020.2011.2002.365361.

9. Brum, J.R., Hurwitz, B.L., Schofield, O., Ducklow, H.W. & Sullivan, M.B. (2016). Seasonal time bombs: dominant temperate viruses affect Southern Ocean microbial dynamics. The ISME Journal, 10, 437–449.

10. Buckling, A. & Rainey, P.B. (2002). The role of parasites in sympatric and allopatric host diversification. Nature, 420, 496–499.

11. Burmeister, A.R., Lenski, R.E. & Meyer, J.R. (2016). Host coevolution alters the adaptive landscape of a virus. Proc Biol Sci, 283.

12. Burmeister, A.R., Sullivan, R.M., Gallie, J. & Lenski, R.E. (2021). Sustained coevolution of phage Lambda and Escherichia coli involves inner-as well as outer-membrane defences and counter-defences. Microbiology, 167.

13. Chaudhry, W.N., Pleška, M., Shah, N.N., Weiss, H., McCall, I.C., Meyer, J.R. et al. (2018). Leaky resistance and the conditions for the existence of lytic bacteriophage. PLOS Biology, 16, e2005971.

14. Childs, L.M., Held, N.L., Young, M.J., Whitaker, R.J. & Weitz, J.S. (2012). Multiscale model of CRISPR-induced coevolutionary dynamics: diversification at the interface of Lamarck and Darwin. Evolution, 66, 2015–2029.

15. Clokie, M.R.J., Millard, A.D., Letarov, A.V. & Heaphy, S. (2011). Phages in nature. Bacteriophage, 1, 31–45.

16. Cooper, V.S., Schneider, D., Blot, M. & Lenski, R.E. (2001). Mechanisms causing rapid and parallel losses of ribose catabolism in evolving populations of Escherichia coli B. Journal of Bacteriology, 183, 2834–2841.

17. Deatherage, D.E. & Barrick, J.E. (2014). Identification of mutations in laboratory-evolved microbes from next-generation sequencing data using breseq. Methods Mol Biol, 1151, 165–188.

18. Erni, B., Zanolari, B. & Kocher, H.P. (1987). The mannose permease of Escherichia coli consists of three different proteins. Amino acid sequence and function in sugar transport, sugar phosphorylation, and penetration of phage lambda DNA. J Biol Chem, 262, 5238–5247.

19. Esquinas-Rychen, M. & Erni, B. (2001). Facilitation of bacteriophage lambda DNA injection by inner membrane proteins of the bacterial phosphoenolpyruvate : carbohydrate phosphotransferase system (PTS). Journal of Molecular Microbiology and Biotechnology, 3, 361–370.

20. Flores, C.O., Meyer, J.R., Valverde, S., Farr, L. & Weitz, J.S. (2011). Statistical-structure of host-phage interactions. Proc Natl Acad Sci U S A, 108, E288–297.

21. Flores, C.O., Poisot, T., Valverde, S. & Weitz, J.S. (2016). BiMat: a MATLAB package to facilitate the analysis of bipartite networks. Methods in Ecology and Evolution, 7, 127–132.

22. Flores, C.O., Valverde, S. & Weitz, J.S. (2013). Multi-scale structure and geographic drivers of cross-infection within marine bacteria and phages. The ISME Journal, 7, 520–532.

23. Fuhrman, J.A. (1999). Marine viruses and their biogeochemical and ecological effects. Nature, 399, 541–548.

24. Gaba, S. & Ebert, D. (2009). Time-shift experiments as a tool to study antagonistic coevolution. Trends Ecol Evol, 24, 226–232.

25. Gandon, S., Buckling, A., Decaestecker, E. & Day, T. (2008). Host-parasite coevolution and patterns of adaptation across time and space. J Evol Biol, 21, 1861–1866.

26. Guimarães, P.R., Jr., Sazima, C., dos Reis, S.F. & Sazima, I. (2007). The nested structure of marine cleaning symbiosis: is it like flowers and bees? Biol Lett, 3, 51–54.

27. Gómez, P. & Buckling, A. (2011). Bacteria-phage antagonistic coevolution in soil. Science, 332, 106–109.

28. Hall, A.R., Scanlan, P.D., Morgan, A.D. & Buckling, A. (2011). Host-parasite coevolutionary arms races give way to fluctuating selection. Ecol Lett, 14, 635–642.

29. Hampton, H.G., Watson, B.N.J. & Fineran, P.C. (2020). The arms race between bacteria and their phage foes. Nature, 577, 327–336.

30. James, A., Pitchford, J.W. & Plank, M.J. (2012). Disentangling nestedness from models of ecological complexity. Nature, 487, 227–230.

31. Katoh, K., Misawa, K., Kuma, K. & Miyata, T. (2002). MAFFT: a novel method for rapid multiple sequence alignment based on fast Fourier transform. Nucleic Acids Res, 30, 3059–3066.

32. Koelle, K., Cobey, S., Grenfell, B. & Pascual, M. (2006). Epochal Evolution Shapes the Phylodynamics of Interpandemic Influenza A (H3N2) in Humans. Science, 314, 1898.

33. Koskella, B. & Brockhurst, M.A. (2014). Bacteria-phage coevolution as a driver of ecological and evolutionary processes in microbial communities. FEMS microbiology reviews, 38, 916–931.

34. Labrie, S.J., Samson, J.E. & Moineau, S. (2010). Bacteriophage resistance mechanisms. Nature Reviews Microbiology, 8, 317–327.

35. Lang, G.I., Rice, D.P., Hickman, M.J., Sodergren, E., Weinstock, G.M., Botstein, D. et al. (2013). Pervasive genetic hitchhiking and clonal interference in forty evolving yeast populations. Nature, 500, 571–574.

36. Leung, C.Y. & Weitz, J.S. (2016). Conflicting attachment and the growth of bipartite networks. Physical Review E, 93, 032303.

37. Luria, S.E. (1945). Mutations of Bacterial Viruses Affecting Their Host Range. Genetics, 30, 84–99.

38. Luria, S.E. & Delbrück, M. (1943). Mutations of Bacteria from Virus Sensitivity to Virus Resistance. Genetics, 28, 491–511.

39. Maddamsetti, R., Johnson, D.T., Spielman, S.J., Petrie, K.L., Marks, D.S. & Meyer, J.R. (2018). Gain-of-function experiments with bacteriophage lambda uncover residues under diversifying selection in nature. Evolution, 72, 2234–2243.

40. Makarenkov, V. (2001). T-REX: reconstructing and visualizing phylogenetic trees and reticulation networks. Bioinformatics, 17, 664–668.

41. Martin, M. (2011). Cutadapt removes adapter sequences from high-throughput sequencing reads. EMBnet. journal, 17, pp. 10–12.

42. Maynard, N.D., Birch, E.W., Sanghvi, J.C., Chen, L., Gutschow, M.V. & Covert, M.W. (2010). A Forward-Genetic Screen and Dynamic Analysis of Lambda Phage Host-Dependencies Reveals an Extensive Interaction Network and a New Anti-Viral Strategy. Plos Genetics, 6, 15.

43. Meyer, J.R., Dobias, D.T., Medina, S.J., Servilio, L., Gupta, A. & Lenski, R.E. (2016). Ecological speciation of bacteriophage lambda in allopatry and sympatry. Science, 354, 1301–1304.

44. Meyer, J.R., Dobias, D.T., Weitz, J.S., Barrick, J.E., Quick, R.T. & Lenski, R.E. (2012). Repeatability and contingency in the evolution of a key innovation in phage lambda. Science, 335, 428–432.

45. Meyer, J.R. & Lenski, R.E. (2020). Subtle Environmental Differences have Cascading Effects on the Ecology and Evolution of a Model Microbial Community*. In: Evolution in Action: Past, Present and Future: A Festschrift in Honor of Erik D. Goodman (eds. Banzhaf, W, Cheng, BHC, Deb, K, Holekamp, KE, Lenski, RE, Ofria, C et al.). Springer International Publishing Cham, pp. 273–288.

46. Petrie, K.L., Palmer, N.D., Johnson, D.T., Medina, S.J., Yan, S.J., Li, V. et al. (2018). Destabilizing mutations encode nongenetic variation that drives evolutionary innovation. Science, 359, 1542–1545.

47. Rodriguez-Valera, F., Martin-Cuadrado, A.-B., Rodriguez-Brito, B., Pašić, L., Thingstad, T.F., Rohwer, F. et al. (2009). Explaining microbial population genomics through phage predation. Nature Reviews Microbiology, 7, 828–836.

48. Sagulenko, P., Puller, V. & Neher, R.A. (2018). TreeTime: Maximum-likelihood phylodynamic analysis. Virus Evol, 4, vex042.

49. Sambrook, J. & Russell, D.W. (2001). Molecular cloning : a laboratory manual. 3rd edn. Cold Spring Harbor Laboratory Press, Cold Spring Harbor, N.Y.

50. Sasaki, A. (2000). Host-parasite coevolution in a multilocus gene-for-gene system. Proc Biol Sci, 267, 2183–2188.

51. Scandella, D. & Arber, W. (1976). Phage λ DNA injection into Escherichia coli pel-mutants is restored by mutations in phage genes V or H. Virology, 69, 206–215.

52. Souza, V., Turner, P.E. & Lenski, R.E. (1997). Long-term experimental evolution in Escherichia coli. V. Effects of recombination with immigrant genotypes on the rate of bacterial evolution. Journal of Evolutionary Biology, 10, 743–769.

53. Stamatakis, A. (2014). RAxML version 8: a tool for phylogenetic analysis and post-analysis of large phylogenies. Bioinformatics, 30, 1312–1313.

54. Stern, A. & Sorek, R. (2011). The phage-host arms race: Shaping the evolution of microbes. BioEssays, 33, 43–51.

55. Suttle, C.A. (2007). Marine viruses--major players in the global ecosystem. Nat Rev Microbiol, 5, 801–812.

56. Thomas, R., Berdjeb, L., Sime-Ngando, T. & Jacquet, S. (2011). Viral abundance, production, decay rates and life strategies (lysogeny versus lysis) in Lake Bourget (France). Environ Microbiol, 13, 616–630.

57. Torsvik, V., Øvreås, L. & Thingstad, T.F. (2002). Prokaryotic diversity--magnitude, dynamics, and controlling factors. Science, 296, 1064–1066.

58. Valverde, S., Elena, S.F. & Solé, R. (2017). Spatially induced nestedness in a neutral model of phage-bacteria networks. Virus Evol, 3, vex021.

59. Volz, E.M., Koelle, K. & Bedford, T. (2013). Viral Phylodynamics. PLOS Computational Biology, 9, e1002947.

60. Weinberger, A.D., Sun, C.L., Pluciński, M.M., Denef, V.J., Thomas, B.C., Horvath, P. et al. (2012). Persisting Viral Sequences Shape Microbial CRISPR-based Immunity. PLOS Computational Biology, 8, e1002475.

61. Weitz, J.S., Hartman, H. & Levin, S.A. (2005). Coevolutionary arms races between bacteria and bacteriophage. Proc Natl Acad Sci U S A, 102, 9535–9540.

62. Weitz, J.S., Poisot, T., Meyer, J.R., Flores, C.O., Valverde, S., Sullivan, M.B. et al. (2013). Phage-bacteria infection networks. Trends Microbiol, 21, 82–91.

63. Weitz, J.S. & Wilhelm, S.W. (2012). Ocean viruses and their effects on microbial communities and biogeochemical cycles. F1000 Biol Rep, 4, 17.

64. Wichman, H.A., Millstein, J. & Bull, J.J. (2005). Adaptive Molecular Evolution for 13,000 Phage Generations. Genetics, 170, 19.

65. Wilhelm, S.W. & Suttle, C.A. (1999). Viruses and Nutrient Cycles in the Sea: Viruses play critical roles in the structure and function of aquatic food webs. BioScience, 49, 781–788.

