## Supplementary Information for "Leapfrog dynamics in phage-bacteria coevolution revealed by joint analysis of cross-infection phenotypes and whole genome sequencing"

#### Δ777 bp IS element deletion's effects on λ resistance

We tested for direct and interactive effects between a *malT*⁻ mutation and the 777 bp deletion by comparing the resistance of strains with all combinations of the two mutations. We already possessed the ancestral strain with no mutations (REL606). Previous to this study we isolated a strain with just the *malT*⁻ (Chaudhry *et al.* 2018). From sequencing isolates from the coevolution experiment we uncovered two strains that possessed these two mutations and no others (isolates D37-7 and D37-9 from day 37). The only strain we did not have was one with the 777 bp deletion. To generate this strain, we selected *malT* revertants of D37-7 and D37-9. The 25-base duplication in *malT* that these strains possess has a high rate of reversion (Chaudhry *et al.* 2018), allowing us to select *malT*⁺ revertants by growing on a medium where maltose is the only carbon source. We spread ~$10^8$ cells of each isolate on minimal maltose agar plates and incubated the plates for two days at 37°C (plate recipes are provided in the next section). A single colony was picked from each plate and re-streaked on tetrazolium maltose (TM) indicator plates to confirm the reversion. Revertants that can metabolize maltose produce white colonies and are easily distinguishable from *malT*⁻ strains that produce smaller red colonies (Shuman & Silhavy 2003). The TM plates were incubated at 37 °C overnight, and this procedure was repeated once more to further purify the isolates. The resulting strains were labeled D37-7+ and D37-9+.

To test for resistance, we compared the growth of the bacteria with and without the phage in wells of a 96-well flat bottom plate. We chose a phage isolate that had a broad host-range to challenge the *E. coli* genotypes (isolate D28-11 from day 28). Six *E. coli* genotypes were evaluated: ancestor without mutations, a strain with only 25-base *malT* duplication, D37-7+, D37-9+, D37-7, and D37-9.

A single plate was used for all trials with LBM9 medium and a total volume of 200 μl . ~100 cells of the bacterial strain and ~$10^9$ λ particles were added to the wells. Four replicates for each host were set up without phage and four with phage. The plates were incubated at 37 °C for about 16 hours in a Tecan Sunrise plate reader. Bacterial densities were observed every 5 minutes by reading optical density (OD) at wavelength 600 nm, just after the plate was shaken.

The growth patterns without the phage were similar for all *E. coli* strains (Fig. S2). None of the strains grew with λ, except for D37-7 and D37-9, the strains with both mutations. To compare the suppressive effect of λ on the different *E. coli* strains, we quantified the maximum OD achieved for each replicate with λ added. We ran a general linear model where maximum OD was predicted by the presence of *malT*⁻, Δ777, and their interactions. A variable that accounts for differences between the strains derived from isolate D37-7 or D37-9 was left out of the model because we found no significant difference between the maximum ODs of D37-7 and D37-9 or D37-7+ and D37-9+. The analysis was performed in R (version 3.6.1). Neither *malT*⁻ or Δ777 conferred resistance: $t$ = -0.098, $P$ = 0.923, and $t$ = 0.045, $P$ = 0.964, respectively. However, the interaction term was highly significant: $t$ = 41.316, $P < 0.001$. This shows that the *malT* mutation and Δ777 bp mutation have an epistatic interaction that confers resistance to λ.

Media recipe for plates

*Minimal maltose agar plates*: 5.34 g potassium phosphate dibasic anhydrous, 2 g potassium phosphate monobasic anhydrous, 1 g ammonium sulfate, 0.57 g sodium citrate dihydrate, 16 g agar, 4 g maltose per liter of water and supplemented to a final concentration of 1 mM magnesium sulfate, 0.0002% w/v thiamine, and 0.0002% w/v biotin.

*Tetrazolium maltose plates*: 10 g tryptone, 1 g yeast extract, 5 g sodium chloride, 16 g agar, 10 g maltose per liter of water and supplemented to a final concentration of 0.005% tetrazolium indicator dye TTC.

**Test of whether phage isolation using REL606 biased phage sampling**

We used the *E. coli* strain (REL606) that was used to initiate the coevolution experiment in order to sample phages. This strategy seemed reasonable since phages sampled from all time points were able to produce plaques on REL606 lawns. However, it is possible that using this standard host may have biased phage sampling if, during coevolution, some phages evolved enhanced infectivity on the coevolved host genotypes and lost (or decreased) their ability to infect REL606. To test this, we re-isolated phages from day 22 using coevolved host genotypes. We chose day 22 because phage genetic and phenotypic diversity peaked at this time point and we reasoned that high diversity samples would be most susceptible to sampling bias. We used 15 coevolved bacteria as well as REL606 as a control. The 15 isolates were from different time points and represent all of the unique genotypes of bacteria that we sampled from the original study. 8 plaques were randomly selected from each host lawn except for three hosts in which no plaques appeared because the bacteria were completely resistant to the day 22 phages. The new isolates were then plated on REL606 to determine if they could infect the ancestral bacteria. Indeed, all 104 new isolates produced clearing on REL606 (Table S3). Next, we performed a more sensitive infection assay to test whether the phages had decreased ability to produce plaques on REL606. We randomly isolated a single phage from each of the different hosts, and then measured the ability of each phage to produce plaques on the host they were isolated on relative to REL606. No phage was less infective on REL606 (Table S3). Altogether, these studies show that REL606 was well-suited to sample the coevolved phage diversity.

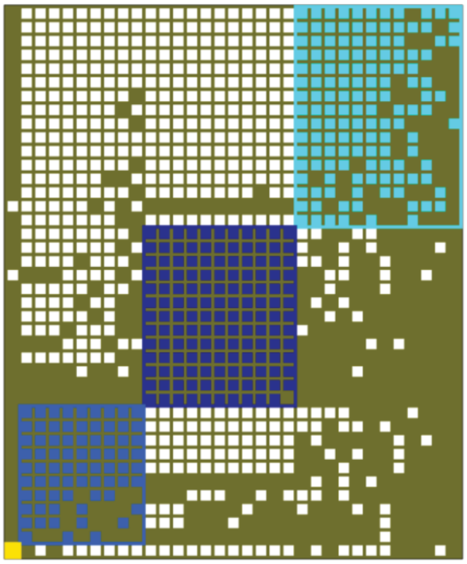

**Fig S1. Phage (columns) and bacterial (rows) interaction network when optimized for modularity.** The original network in Fig. 1A but reassembled to calculate modularity using the software BiMat. Filled squares indicate a combination of host and phage that result in successful interactions (EOP > 0). The measured value of modularity ($Q_b$) was 0.0983 vs. the null value of 0.0884 ± 0.0038 based on 200 trials.

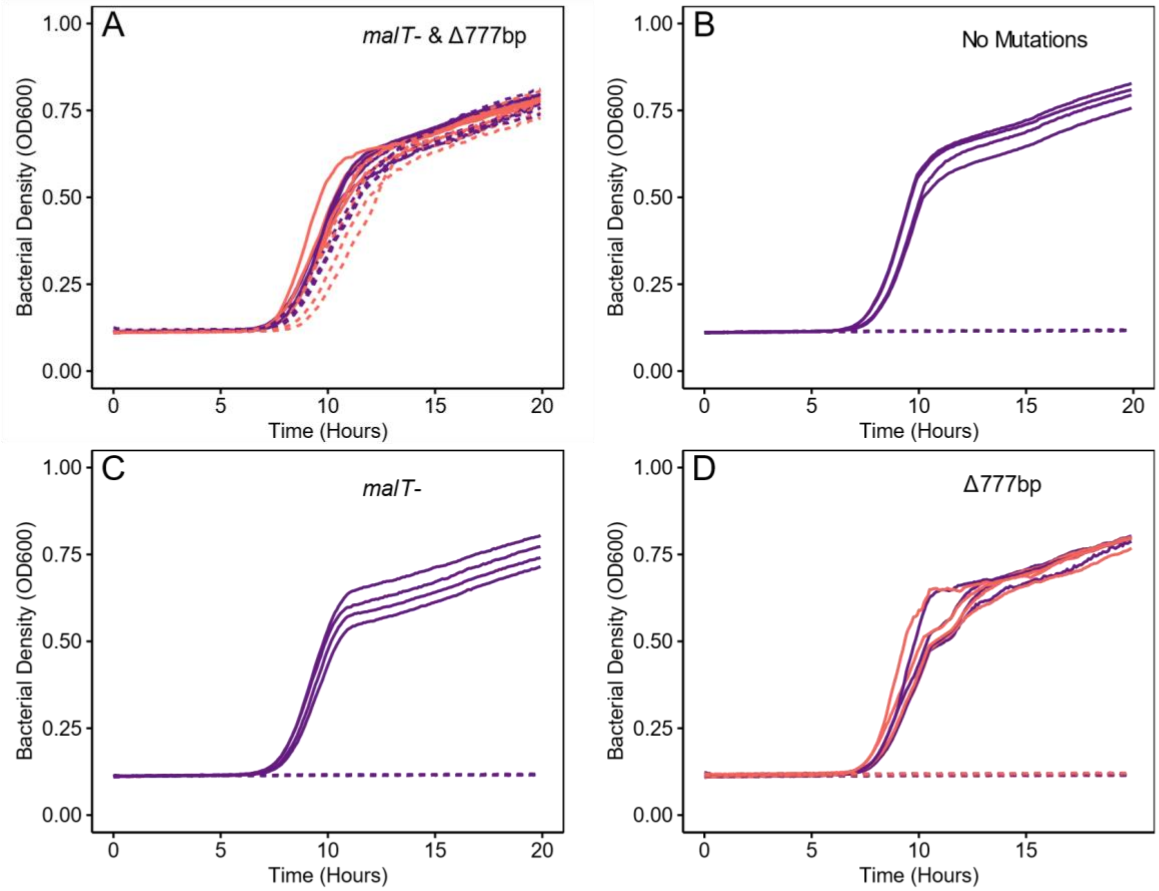


**Fig. S2.** Growth trajectories of different bacterial strains growing in the absence (solid line) or presence (dashed line) of a coevolved phage isolated from day 28 in wells of a 96-well plate. **A)** Day 37 isolates D37-7 (purple) and D37-9 (pink), which have both *malT* and Δ777 mutations, **B)** ancestor REL606 which has neither *malT* nor Δ777 mutations, **C)** *malT*— strain which only has the 25-base duplication *malT* mutation, and **D)** strains D37-7+ and D37-9+ derived from D37-7 and and D37-9 (colors as in Panel A) which have reverted their *malT* mutation from MalT— to $MalT^+$, and thus, have only the Δ777 mutation.

**Table S1**. Genomic variation present in *E. coli* population on Day 8. The mutations are annotated with respect to the ancestral genome (GenBank: CP000819.1). The mutations marked in red have been shown to confer resistance to λ. The mutation in the bold indicates the dominant lineage (marked in red in Figure 3) at the end of coevolution.

| Genome Location | Mutation | Gene | Product |
|---|---|---|---|
| 38,192 | G→T | *carB* → / → *caiF* | carbamoyl-phosphate synthase large subunit/DNA-binding transcriptional activator |
| 38,193 | C→T | *carB* → / → *caiF* | carbamoyl-phosphate synthase large subunit/DNA-binding transcriptional activator |
| 38,194 | C→T | *carB* → / → *caiF* | carbamoyl-phosphate synthase large subunit/DNA-binding transcriptional activator |
| 38,195 | C→T | *carB* → / → *caiF* | carbamoyl-phosphate synthase large subunit/DNA-binding transcriptional activator |
| 38,196 | A→T | *carB* → / → *caiF* | carbamoyl-phosphate synthase large subunit/DNA-binding transcriptional activator |
| 38,199 | A→T | *carB* → / → *caiF* | carbamoyl-phosphate synthase large subunit/DNA-binding transcriptional activator |
| 38,200 | A→T | *carB* → / → *caiF* | carbamoyl-phosphate synthase large subunit/DNA-binding transcriptional activator |
| 386,921 | C→T | *phoR* → | sensory histidine kinase in two-component regulatory system with PhoB |
| 519,803 | G→T | *fdrA* → | membrane protein FdrA |
| 519,808 | A→C | *fdrA* → | membrane protein FdrA |
| 560,154 | C→T | *ompT* ← | outer membrane protease VII (outer membrane protein 3b) |
| 863,867 | G→A | *yliC* → | predicted peptide transporter subunit: membrane component of ABC superfamily |
| 863,868 | G→T | *yliC* → | predicted peptide transporter subunit: membrane component of ABC superfamily |
| 863,873 | A→C | *yliC* → | predicted peptide transporter subunit: membrane component of ABC superfamily |
| 863,874 | T→C | *yliC* → | predicted peptide transporter subunit: membrane component of ABC superfamily |
| 949,387 | G→A | *trxB* ← / → *lrp* | thioredoxin reductase, FAD/NAD(P)-binding/DNA-binding transcriptional dual regulator, leucine-binding |
| 1,368,412:1 | $(T)_{9\rightarrow10}$ | *pspE* → / → *ycjM* | thiosulfate:cyanide sulfurtransferase (rhodanese)/predicted glucosyltransferase |
| 1,418,284 | G→T | *rzpR* → | predicted defective peptidase |
| 1,605,635 | Δ1 bp | *stfR* ← | predicted tail fiber protein |
| 1,605,636 | T→G | *stfR* ← | predicted tail fiber protein |
| 1,605,637 | G→T | *stfR* ← | predicted tail fiber protein |
| 1,605,637:1 | +T | *stfR* ← | predicted tail fiber protein |
| 1,881,837 | Δ1 bp | *manY* → | mannose-specific enzyme IIC component of PTS |
| 1,881,838 | Δ1 bp | *manY* → | mannose-specific enzyme IIC component of PTS |
| 1,882,021 | C→T | *manY* → | mannose-specific enzyme IIC component of PTS |

| | | | |
|---|---|---|---|
| 1,882,908:1 | +A | *manZ* → | mannose-specific enzyme IID component of PTS |
| **1,882,915** | **Δ16 bp** | ***manZ* →** | **mannose-specific enzyme IID component of PTS** |
| 2,111,270 | C→A | *ECB_01999* → | putative phage protein |
| 2,250,122:1 | +G | *napG* ← | quinol dehydrogenase periplasmic component |
| 2,250,126 | A→C | *napG* ← | quinol dehydrogenase periplasmic component |
| 2,250,129 | Δ1 bp | *napG* ← | quinol dehydrogenase periplasmic component |
| 2,310,865 | G→A | *yfaZ* ← / → *yfaO* | predicted outer membrane porin protein/predicted NUDIX hydrolase |
| 2,310,868 | C→A | *yfaZ* ← / → *yfaO* | predicted outer membrane porin protein/predicted NUDIX hydrolase |
| 2,401,525 | Δ1 bp | *yfcX* ← | fused enoyl-CoA hydratase and epimerase and isomerase/3-hydroxyacyl-CoA dehydrogenase |
| 2,401,526 | G→A | *yfcX* ← | fused enoyl-CoA hydratase and epimerase and isomerase/3-hydroxyacyl-CoA dehydrogenase |
| 2,401,527 | C→A | *yfcX* ← | fused enoyl-CoA hydratase and epimerase and isomerase/3-hydroxyacyl-CoA dehydrogenase |
| 2,401,529 | A→T | *yfcX* ← | fused enoyl-CoA hydratase and epimerase and isomerase/3-hydroxyacyl-CoA dehydrogenase |
| 2,940,619 | A→T | *ygfB* ← | hypothetical protein |
| 3,000,508 | C→G | *flu* → | antigen 43 (Ag43) phase-variable biofilm formation autotransporter |
| 3,482,706:1 | 25-bp duplication | *malT* → | transcriptional regulator MalT |
| **3,482,802** | **Δ141 bp** | ***malT* →** | **transcriptional regulator MalT** |
| 3,483,094:1 | +C | *malT* → | transcriptional regulator MalT |
| 3,483,094:2 | +T | *malT* → | transcriptional regulator MalT |
| 3,942,902 | T→A | *yifK* → | predicted transporter |
| 4,236,155 | T→A | *lexA* → | LexA repressor |
| 4,236,156 | T→A | *lexA* → | LexA repressor |
| 4,236,158 | C→G | *lexA* → | LexA repressor |
| 4,236,160 | T→A | *lexA* → | LexA repressor |
| 4,236,161 | T→A | *lexA* → | LexA repressor |
| 4,300,483 | C→T | *phnG* ← | carbon-phosphorus lyase complex subunit |
| 4,504,878 | T→A | *insA-25* → / → *ECB_04162* | IS1 protein InsA/hypothetical protein |
| 4,537,685 | A→T | *yjiC* ← / → *yjiD* | hypothetical protein/DNA replication/recombination/repair protein |

**Table S2**. Genomic variation present in the phage population on Day 8 of the coevolution experiment as compared to the ancestral λ strain cI26 used in the study. The location of the mutations is annotated with respect to the λ reference genome (GenBank: NC_001416). The mutations which have fixed in the population are italicized. The mutation in the bold indicates the dominant lineage (marked in blue in Figure 3) at the end of coevolution.

| **Genome Location** | **Mutation** | **Amino acid change** | **Gene** | **Product** |
|---|---|---|---|---|
| 11,445 | C→T | A->V | *H* → | Tail component |
| **11,451** | **C→T** | **A->V** | ***H*** **→** | **Tail component** |
| 15,890 | A→G | D->G | *J* → | Tail- host specificity protein |
| 16,218 | G→T | | *J* → | Tail- host specificity protein |
| 16,227 | T→C | | *J* → | Tail- host specificity protein |
| 16,299 | A→G | | *J* → | Tail- host specificity protein |
| 16,318 | A→C | M->L | *J* → | Tail- host specificity protein |
| 16,319 | T→C | M->T | *J* → | Tail- host specificity protein |
| 16,350 | T→C | | *J* → | Tail- host specificity protein |
| 16,449 | C→T | | *J* → | Tail- host specificity protein |
| 16,485 | G→C | | *J* → | Tail- host specificity protein |
| 16,497 | A→G | | *J* → | Tail- host specificity protein |
| 16,524 | C→T | | *J* → | Tail- host specificity protein |
| 16,596 | G→A | | *J* → | Tail- host specificity protein |
| 16,599 | G→A | | *J* → | Tail- host specificity protein |
| 16,606 | A→G | T->A | *J* → | Tail- host specificity protein |
| 16,607 | C→T | T->M | *J* → | Tail- host specificity protein |
| 16,725 | C→T | | *J* → | Tail- host specificity protein |
| 16,774 | G→C | A->P | *J* → | Tail- host specificity protein |
| 16,775 | C→T | A->V | *J* → | Tail- host specificity protein |
| 16,791 | T→C | | *J* → | Tail- host specificity protein |
| 16,794 | T→C | | *J* → | Tail- host specificity protein |
| 16,866 | A→G | | *J* → | Tail- host specificity protein |
| 16,869 | A→G | | *J* → | Tail- host specificity protein |
| 16,893 | T→C | | *J* → | Tail- host specificity protein |
| 16,902 | C→G | | *J* → | Tail- host specificity protein |
| 16,905 | C→T | | *J* → | Tail- host specificity protein |
| 16,908 | A→C | | *J* → | Tail- host specificity protein |
| 16,938 | T→C | | *J* → | Tail- host specificity protein |
| 16,972 | A→C | S->R | *J* → | Tail- host specificity protein |
| 16,980 | T→C | | *J* → | Tail- host specificity protein |
| 16,983 | T→G | | *J* → | Tail- host specificity protein |
| 16,986 | T→C | | *J* → | Tail- host specificity protein |
| 16,998 | G→A | | *J* → | Tail- host specificity protein |
| *18,503* | *C→T* | *A->V* | *J* → | *Tail- host specificity protein* |
| *18,734* | *T→C* | *V->A* | *J* → | *Tail- host specificity protein* |
| *18,823* | *G→A* | *D->N* | *J* → | *Tail- host specificity protein* |
| *18,868* | *A→T* | *I->F* | *J* → | *Tail- host specificity protein* |

**Table S3: Results from the test of possible phage isolation bias**. EOP is the number of plaques produced on the isolation host divided by plaques produced on REL606. Error in this assay was estimated for the REL606 control by repeating the measurement four separate times. The 95% confidence interval is reported in the table.

| Host genotype ID used to isolate λ | Proportion of isolates able to infect REL606 | EOP |
|---|---|---|
| **D_8_1** | 8 of 8 | 0.880 |
| **D_8_2** | 8 of 8 | 0.368 |
| **D_8_3** | 8 of 8 | 0.222 |
| **D_8_4** | 8 of 8 | 0.333 |
| **D_8_6** | 8 of 8 | 0.133 |
| **D_8_7** | 8 of 8 | 0.600 |
| **D_15_2** | 8 of 8 | 0.938 |
| **D_15_3** | 8 of 8 | 0.267 |
| **D_15_7** | 8 of 8 | 1.000 |
| **D_22_1** | 8 of 8 | 0.714 |
| **D_28_4** | 8 of 8 | 0.560 |
| **D_28_6** | NA (completely resistant genotype) | - |
| **D_28_8** | NA (completely resistant genotype) | - |
| **D_28_10** | 8 of 8 | 1.375 |
| **D_37_2** | NA (completely resistant genotype) | - |
| **REL606 (control)** | 8 of 8 | 1.020 ± 0.447 |

### References


1.
Chaudhry, W.N., Pleška, M., Shah, N.N., Weiss, H., McCall, I.C., Meyer, J.R. *et al.* (2018). Leaky resistance and the conditions for the existence of lytic bacteriophage. *PLOS Biology*, 16, e2005971.

2.
Shuman, H.A. & Silhavy, T.J. (2003). The art and design of genetic screens: Escherichia coli. *Nat Rev Genet*, 4, 419-431.
